# Previously reward-associated stimuli capture spatial attention in the absence of changes in the corresponding sensory representations as measured with MEG

**DOI:** 10.1101/622589

**Authors:** L Tankelevitch, E Spaak, MFS Rushworth, MG Stokes

**Affiliations:** Department of Experimental Psychology, Wellcome Centre for Integrative Neuroimaging, University of Oxford, Oxford, OX1 3UD, United Kingdom; Oxford Centre for Human Brain Activity, Wellcome Centre for Integrative Neuroimaging, Department of Psychiatry, University of Oxford, Oxford, OX3 7JX, United Kingdom; Centre for Functional MRI of the Brain, Wellcome Centre for Integrative Neuroimaging, Nuffield Department of Clinical Neurosciences, John Radcliffe Hospital, University of Oxford, Oxford, OX3 9DU, United Kingdom; Donders Institute for Brain, Cognition and Behaviour, Radboud University Nijmegen, 6500 HB Nijmegen, The Netherlands

## Abstract

Studies of selective attention typically consider the role of task goals or physical salience, but recent work has shown that attention can also be captured by previously reward-associated stimuli, even if they are currently task-irrelevant. One theory underlying this value-driven attentional capture (VDAC) is that reward-associated stimulus representations may undergo plasticity in sensory cortex, thereby automatically capturing attention during early processing. To test this, we used magnetoencephalography to probe whether stimulus location and identity representations in sensory cortex are modulated by reward learning. We furthermore investigated the time-course of these neural effects, and their relationship to behavioural VDAC. Male and female human participants first learned stimulus-reward associations. Next, we measured VDAC in a separate task by presenting these stimuli in the absence of reward contingency, and probing their effects on the processing of separate target stimuli presented at different time lags. Using time-resolved multivariate pattern analysis, we found that learned value modulated the spatial selection of previously rewarded stimuli in posterior visual and parietal cortex from ∼260ms after stimulus onset. This value modulation was related to the strength of participants’ behavioural VDAC effect and persisted into subsequent target processing. Furthermore, we found a spatially invariant value signal from ∼340ms. Importantly, learned value did not influence cortical signatures of early processing (i.e., earlier than ∼200ms), nor did it influence the decodability of the identity of previously rewarded stimuli. Our results suggest that VDAC is underpinned by learned value signals which modulate spatial selection throughout posterior visual and parietal cortex. We further suggest that VDAC can occur in the absence of changes in early visual processing in cortex.

**Significance statement:** Attention is our ability to focus on relevant information at the expense of irrelevant information. It can be affected by previously learned but currently irrelevant stimulus-reward associations, a phenomenon termed “value-driven attentional capture” (VDAC). The neural mechanisms underlying VDAC remain unclear. It has been speculated that reward learning induces visual cortical plasticity which modulates early visual processing to capture attention. Although we find that learned value modulates spatial signals in visual cortical areas, an effect which correlates with VDAC, we find no relevant signatures of changes in early visual processing in cortex.

## Introduction

Factors influencing selective attention have been traditionally categorised as either stemming from task demands (e.g., a red book will capture attention if one is searching for red books) or from physical salience (e.g., a red book among blue books will automatically capture attention) (Beck and Kastner, 2009; Desimone and Duncan, 1995; Egeth and Yantis, 1997). Recently, reward history—e.g., the learned association of a red logo with a rewarding experience—has also been shown to play an important role in shaping attention (Anderson, 2013; Awh et al., 2012; Chelazzi et al., 2013). This has been demonstrated in experiments using an initial “training phase” in which visual stimuli are associated with different reward outcomes. During a later “testing phase”, participants complete a new, formally unrelated task, which re-uses the previously reward-associated stimuli (e.g., Anderson et al., 2011). Critically, these stimuli continue to capture attention even in the absence of any reward contingency, physical salience, or task relevance, a phenomenon termed value-driven attentional capture (VDAC; Anderson et al., 2011; Chelazzi et al., 2014; Raymond and O’Brien, 2009; Theeuwes and Belopolsky, 2012). VDAC is a prime example of how attention is influenced by prior experience; understanding its neural mechanisms is therefore instrumental to bridging the fields of learning and attention (Anderson, 2016; Failing and Theeuwes, 2018; Le Pelley et al., 2016).

The behavioural effects of VDAC imply learning-related plasticity, but the nature of these changes remains unclear (Anderson, 2016; Chelazzi et al., 2013; Failing and Theeuwes, 2018). Some have suggested that stimulus feature representations in sensory cortex may undergo reward-mediated plasticity during learning (Anderson, 2016; Failing and Theeuwes, 2018; Hickey and Peelen, 2017; van Koningsbruggen et al., 2016). Strengthened cortical representations could then rapidly capture attention during early visual processing, potentially leading to the behavioural effects of VDAC (Anderson, 2016; Failing and Theeuwes, 2018). However, evidence for the existence of this mechanism is limited (Anderson, 2016; van Koningsbruggen et al., 2016).

Electroencephalography (EEG) studies of VDAC show various types of value modulations in visual processing (Failing and Theeuwes, 2018). The timing of these effects matters (Luck et al., 2000). Early effects may indicate changes in how reward-associated stimuli are initially processed in sensory cortex, consistent with the visual cortical plasticity account (Anderson, 2016; Failing and Theeuwes, 2018; van Koningsbruggen et al., 2016). In contrast, later effects may reflect attentional modulation triggered by these stimuli, consistent with learned value signals originating elsewhere (Buffalo et al., 2010; Foxe and Simpson, 2002; Itthipuripat et al., 2019). Hickey et al. (2010) and Luque et al. (2017) both report an enhancement of the early sensory-evoked P1 component for reward-associated stimuli, suggesting that value modulates early visual processing of these stimuli. In contrast, Qi et al. (2013) report that reward-associated stimuli only evoked the later N2pc component, reflecting an attentional shift towards these stimuli. Others find that reward-associated distracters affect processing of *competing* stimuli, reducing the P1, N1, and N2pc (Itthipuripat et al., 2015; MacLean and Giesbrecht, 2015a). Thus, existing evidence on the timing of value effects during VDAC is mixed.

Comparison between studies is additionally challenging because in some studies the task-relevance, and therefore the reward history, of stimuli changes across trials (e.g., Hickey et al., 2010; Itthipuripat et al., 2015). In other studies, reward-associated stimuli are irrelevant and unrewarded for the entire task (e.g., MacLean and Giesbrecht, 2015; Qi et al., 2013).

Furthermore, it is important to isolate effects related to the processing of reward-associated stimuli from those related to the processing of competing target stimuli (e.g., Hickey and Peelen, 2015; Itthipuripat et al., 2019).

Finally, most prior studies have examined the *spatial* discriminability of reward-associated stimuli, for example, by testing for changes in the lateralised P1 and N2pc components (e.g., Hickey et al., 2010). However, plasticity-induced strengthening of stimulus representations could also manifest in altered neural discriminability of stimulus *identity* (Hickey and Peelen, 2015; LeMessurier and Feldman, 2018; Poort et al., 2015). This has not been demonstrated in the context of VDAC using time-resolved methods. Hickey and Peelen (2015) used multivariate pattern analysis (MVPA) to show that fMRI representations of previously reward-associated stimuli are indeed modulated in visual cortex, although the use of fMRI precluded insight into the timing of these effects.

In the current study, we used time-resolved MVPA of magnetoencephalography (MEG) data to probe stimulus location and identity representations in sensory cortex during VDAC. After participants learned stimulus-reward associations, we measured VDAC in a separate task by presenting these stimuli in the absence of reward contingency, and probing their effects on processing of separate target stimuli. Critically, we included an interval between previously reward-associated stimuli and the imperative targets to isolate the neural processes triggered by the reward stimuli, uncontaminated by target processing. Consistent with VDAC, we found that learned value modulated the spatial selection of previously rewarded stimuli in posterior visual and parietal cortex. However, these effects only emerged from ∼260ms after stimulus onset, in contrast to previous evidence of early effects. Moreover, learned value did not modulate identity representations of the previously reward-associated stimuli in cortex.

## Materials and methods

### Participants

All experiments were conducted under ethical approval from the Central University Research Ethics Committee of the University of Oxford and with informed consent from participants. We aimed for a sample of 30 human participants from the Oxford community for this experiment. Due to poor behavioral performance or MEG acquisition artefacts, we recruited ‘replacement’ participants as needed, leading to a total sample of 37 participants, aged 19-34 years (M = 24.6 years, SD = 4.2 years; 18 female). All participants were right-handed and had normal or corrected-to-normal vision. One participant was excluded during MEG data acquisition due to inability to follow task instructions, and six participants were excluded prior to data analysis due to artefacts in the MEG data, leaving 30 participants for analysis.

### Study structure

Participants completed a *reward learning task* to induce learning of stimulus-reward associations (training phase), and a visual discrimination task (i.e., *attention task)* to test the effect of these associations on visual attention (testing phase). To maximize learning and increase task engagement, participants completed two sessions of each task in an interleaved order on the same day. Participants were compensated based on their winnings in the reward learning task and were explicitly told that the attention task was not rewarded and was simply a requirement to complete as part of the experiment.

### Reward learning task (training phase)

The task was presented using MATLAB (MathWorks, MA, USA) and Psychtoolbox-3 (Kleiner et al., 2007). Participants sat 120 cm from the display, which was a Panasonic PT D7700E (Osaka, Japan) translucent back-projection display, 54.5 × 43cm, resolution 1280 × 1024px, 60 Hz refresh rate. After a 500-1000ms (randomly jittered) inter-trial interval (ITI), two option stimuli (each sized 2.4 × 2.4° visual angle) appeared on screen (Figure 1a-b). Participants were instructed to maintain fixation until the fixation cross disappeared after 500ms. After the fixation cross offset, participants had 5000ms to indicate their choice with a left or right button press. After the choice, a feedback display presented the outcomes of the chosen and unchosen stimuli. To facilitate learning, the background brightness as well as a presented sound effect indicated the outcome reward magnitude. Participants began the next trial with a button press.

**Figure 1.**
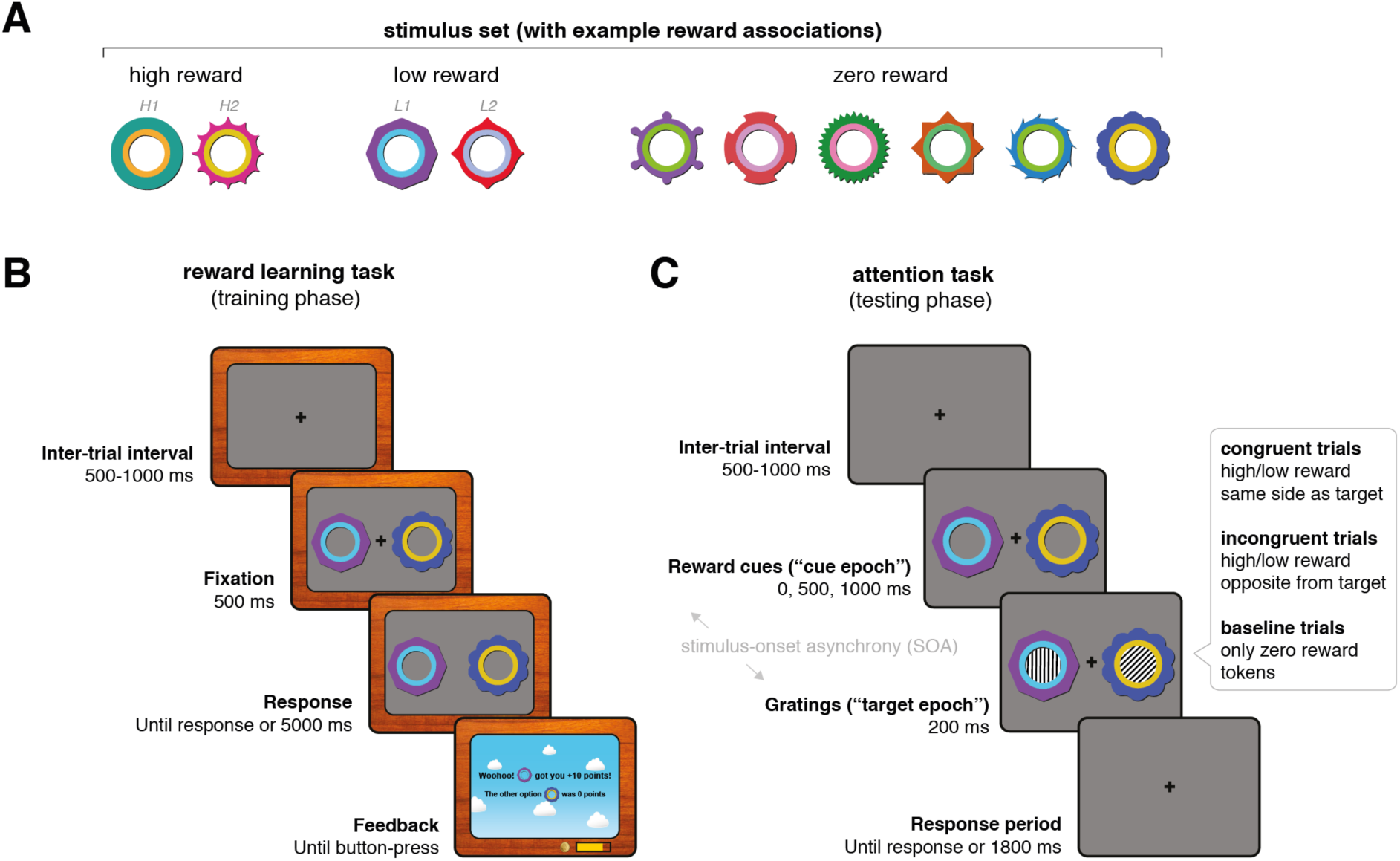
Experimental task design. **(a)** Stimulus set with example associated rewards. Stimulus-reward contingencies were counterbalanced across participants. H1, H2, L1, and L2 refer to reward stimulus labels in the decoding schematics in Figure 3. **(b)** Experimental design for the reward learning task (training phase). Each trial began with a fixation period (500-1000ms, randomly sampled), followed by the presentation of two option stimuli. After 500ms, the fixation cross disappeared and participants were free to indicate their decision by button-press within the time limit (5000ms). Decisions were followed by a feedback display indicating the points obtained for the chosen stimulus, as well as the points that would have been awarded for the unchosen stimulus. Participants initiated the next trial by button-press. Participants could track their earned points and money with on-screen counters. **(c)** Experimental design for the attention task (testing phase). Each trial began with a fixation period (500-1000ms, randomly sampled), followed by the presentation of two reward stimuli from the previous task (now uninformative and unrewarded). Each trial included one high- or low-reward stimulus together with a zero-reward stimulus, or two zero-reward stimuli (baseline trials). After a stimulus-onset asynchrony (0, 500, or 1000ms), gratings appeared inside the reward stimuli for 200ms. One grating was always either vertically or horizontally oriented (the target), and the other was always oriented obliquely, at one of twelve angles (the distracter). In congruent (incongruent) trials, the target grating appeared on the same (opposite) side as the high- or low-reward stimulus. In trials with two zero-reward stimuli, the target could appear on either side. Reward stimuli and gratings disappeared together after 200ms. Participants had an additional 1800ms to indicate with a button-press whether the target is oriented vertically or horizontally. Note that task stimuli are enlarged for illustration purposes.

There were 10 option stimuli that participants learned about across trials: two high reward (250 points), two low reward (25 points), and six zero reward stimuli (0 points; Figure 1a). 1000 points were equivalent to £1, with participants receiving their winnings at the end of the experiment. A ‘points bar’ at the bottom of the screen indicated how many points they had accumulated for the next £1 and a pound (£) counter indicated the total money won thus far. Reward associations were deterministic. To control for possible stimulus difference between option stimuli, reward associations were counterbalanced across participants using a Latin square design. There were 96 High vs. Low reward, 96 High vs. Zero reward, and 192 Low vs. Zero reward choice trials per session, with the latter trial type doubled to equate selection frequency between high and low reward stimuli. Option stimuli varied in both colors and shapes, so participants could use either feature or a combination for learning. Trial order was randomized once and presented consistently across participants to minimize differences in learning.

### Attention task (testing phase)

The hardware and software setup for stimulus presentation was identical to the reward learning task. The task began with a 500-1000ms randomly jittered ITI, during which participants had to fixate on a central cross (Figure 1c). Two of the previously rewarded but currently task-irrelevant reward stimuli then appeared on screen. These could be a pair of High and Zero reward stimuli *(high reward trials)*, Low and Zero reward stimuli (*low reward trials*), or two Zero reward stimuli (*baseline trials*). Following a randomly selected delay of 0, 500, or 1000ms (the *stimulus-onset asynchrony, SOA*), two task-relevant oriented gratings (diameter 1.2° visual angle, with the inner edge positioned 5.4° away from center) appeared inside each of the reward stimuli. The target grating was oriented either 0° or 90°. The participants’ task was to indicate this orientation with a button-press. The non-target (distracter) grating was oriented obliquely at 20-70° or 110-160° degrees, randomly selected from 10° bins. Gratings were presented for 200ms, after which they offset together with the reward stimuli. Participants had up to 1800ms after grating offset to make their response. Auditory feedback was provided with a high frequency tone signifying correct trials and a low frequency tone signifying incorrect trials.

Critically, in high and low reward trials, the target grating appeared inside either the High or Low reward stimuli (*congruent trials*), or inside the Zero reward stimulus (*incongruent trials*). In baseline trials, the target grating could appear in either Zero reward stimulus. Reward stimuli and target locations varied randomly between left and right of the fixation cross.

In each session, there were 60 trials per cell in the reward (3) × congruency (2) × SOA (3) condition matrix, with the exception of the baseline condition in which congruency was not a factor. There were 1800 trials in total across both sessions. All trials were fully randomized across participants. Each session was divided into 19 blocks, with 50 trials per block, and each block taking ∼2 minutes to complete. Participants were instructed to maintain fixation and minimize blinking throughout the duration of a block but were able to take a short break after each block.

In addition to using the SOA as a tool to isolate the processing of reward stimuli, some previous research suggested that VDAC may decrease with increasing SOA duration (e.g., Le Pelley et al., 2013). We manipulated the SOA duration in the current experiment to test for this possibility. However, we found no evidence for an effect of SOA on attentional capture at the behavioural level (in reaction time, our *a priori* measure of interest; see Results and Figure 4a). The absence of an SOA effect is consistent with a pilot behavioural study that we conducted (not reported here), and with some previous research (Failing and Theeuwes, 2015). We also found no evidence of neural differences between the 500ms and 1000ms SOA conditions (not reported here). Therefore, we collapse across these two conditions for all neural analyses (see below for decoding methods).

**Figure 2.**
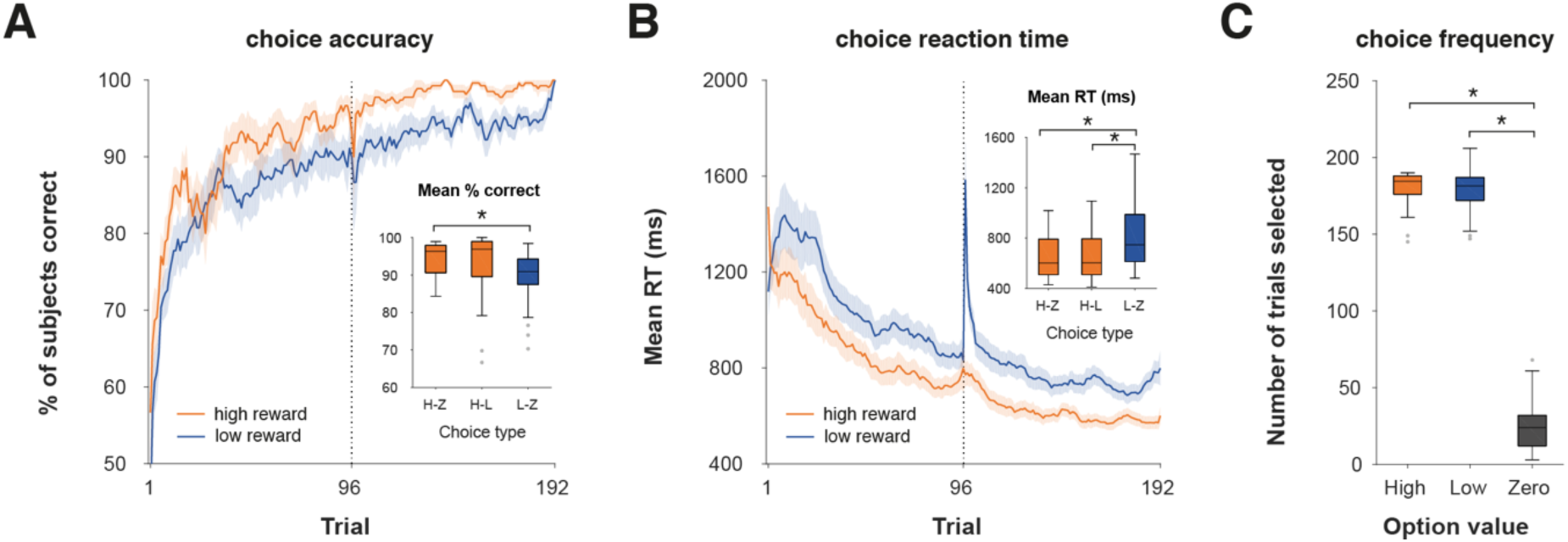
Behavioural results in the reward learning task (training phase). **(a)** Grand-average learning curves for high-reward (*High vs. Zero, High vs. Low*) and low-reward (*Low vs. Zero*) choice trials. Dotted line indicates the boundary between sessions one and two. Error shading reflects standard error of the mean (SEM). *Inset:* Average percent correct as a function of choice type. H-Z = High vs. Zero; H-L = High vs. Low; L-Z = Low vs. Zero. The central mark within each box indicates the median, the box edges indicate the 25^th^ and 75^th^ percentiles, the whiskers indicate the most extreme data points not considered outliers, and the grey points indicate outliers. Asterisks indicate p < 0.05 in a paired t-test. **(b)** Grand-average trial-wise reaction time. Same conventions as (a). *Inset*: Average reaction time as a function of choice type. Same convention as (a). **(c)** Selection frequency as a function of stimulus value. Same conventions as (a).

**Figure 3.**
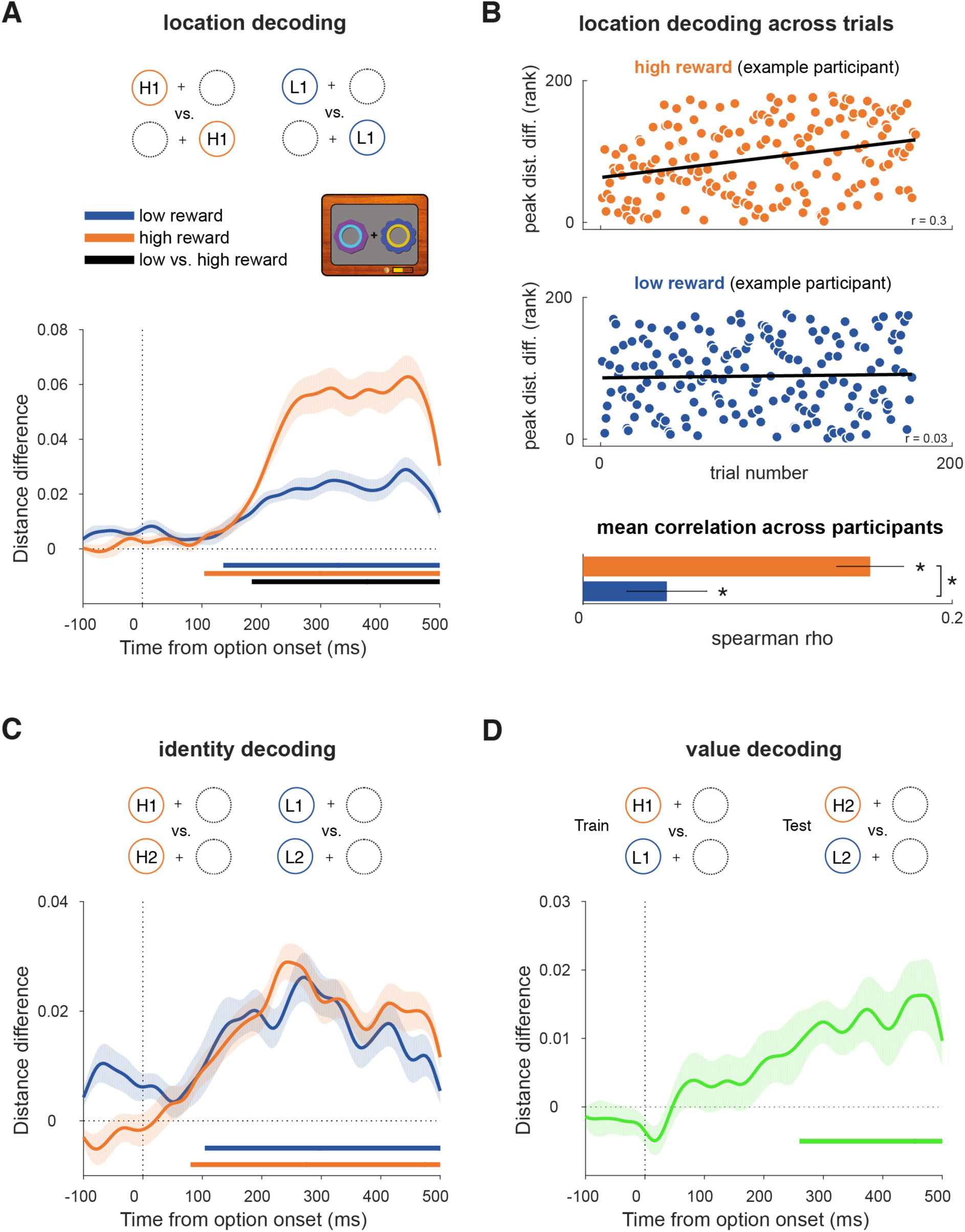
Decoding reward option location, identity, and value in the reward learning task. **(a)** Decoding the location of the reward-associated stimuli involved discriminating between trials in which the higher reward stimulus appeared on the left versus right, collapsing across stimulus identities. Curves depict decoding averaged across trials and participants for high and low reward options. Bars indicate significant decoding (p < 0.05, cluster-based permutation test across time) for each reward condition (orange, blue bars) and between reward conditions (black bar). Shaded regions indicate SEM. **(b)** Correlation between location decoding at peak and trial number in the high and low reward conditions. Scatter plots show ranked peak decoding across trials for one example participant, together with the corresponding trend lines. Bar plot shows mean Fisher z-transformed Spearman correlation for each condition, across all participants. **(c)** Decoding the identity of reward-associated stimuli involved discriminating between trials with distinct stimulus identities for each reward value, within each location with results averaged across (only one location depicted in schematic). Same conventions as Figure 3a. **(d)** Decoding reward value involved using one pair of high and low reward stimuli as training data and a different pair as testing data (collapsing across location); any signal that generalises across pairs is independent of stimulus identity or location and can therefore be attributed to the difference in value between high and low reward stimuli. Curve depicts decoding averaged across trials and participants. Bar indicates significant decoding (p < 0.05, cluster-based permutation test across time). Shaded regions indicate SEM.

**Figure 4.**
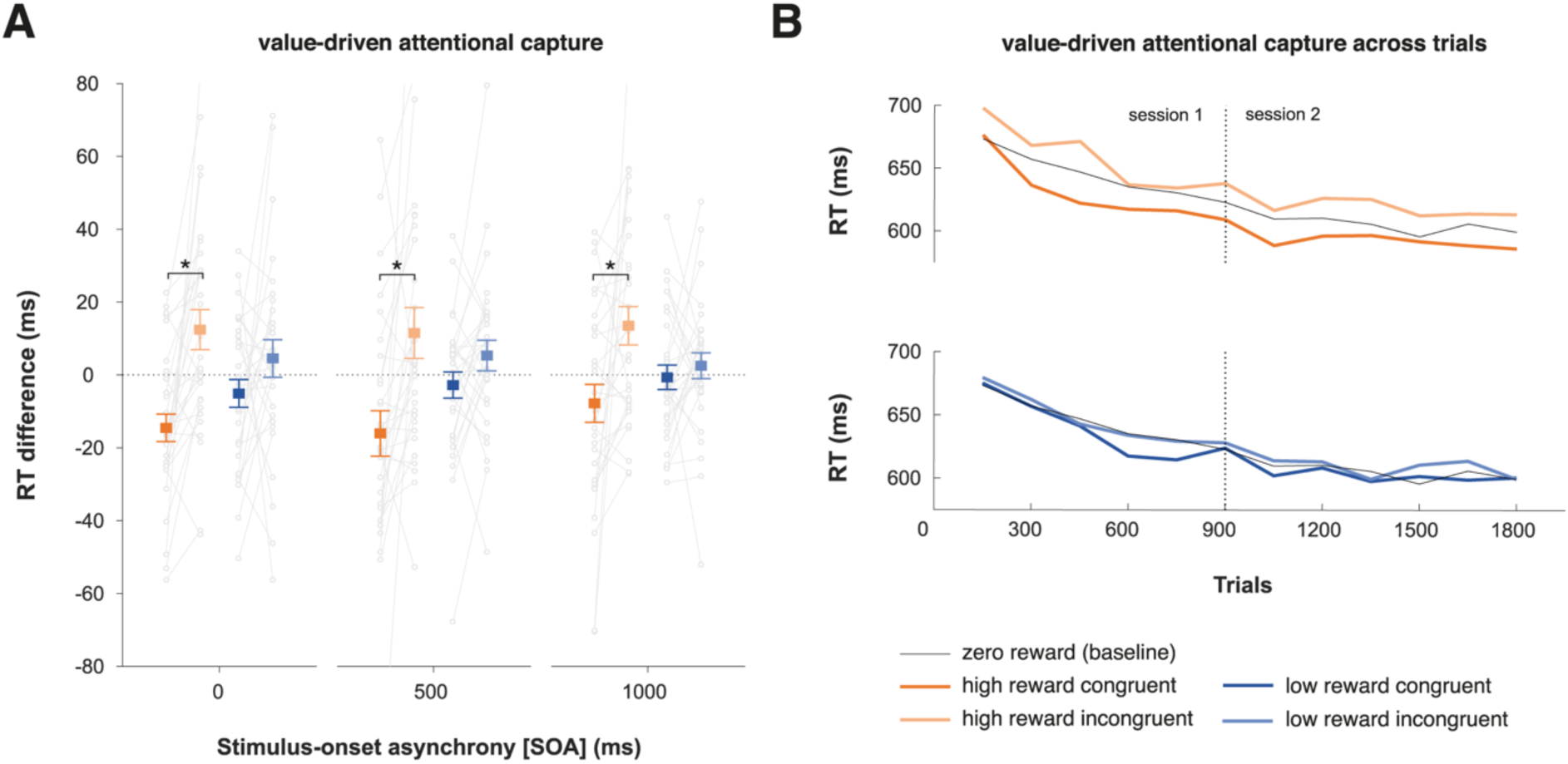
Value-driven attentional capture as measured by behaviour in the attention task. **(a)** Mean RT difference scores (with baseline condition subtracted out) as a function of reward, congruency, and SOA. Grey lines indicate individual participant data. Positive and negative values indicate RTs slower and faster than baseline, respectively. Asterisks indicate p < 0.05 in a paired t-test. Error bars indicate SEM. **(b)** RTs for each condition across trials, averaged within 150-trial bins.

### MEG data acquisition and preprocessing

For both tasks, whole-head MEG data was recorded in a magnetically shielded room using a 306-channel Elekta Neuromag VectorView scanner (102 magnetometers, 204 gradiometers; Elekta, Stockholm, Sweden) at the Oxford Centre for Human Brain Activity (OHBA). Four head position indicator (HPI) coils were placed behind the ears and on top of the forehead to track head movements in the scanner. A magnetic Polhemus Fastrak 3D digitiser pen (Vermont, USA) was used to register the HPI coils and three anatomical landmarks (nasion, left and right auricular points), and digitise the head shape by marking 300 points along the scalp. MEG data were sampled at 1000 Hz with a 0.03-330 Hz bandpass filter during signal digitisation. Electrocardiography (ECG) and electrooculography (EOG) were recorded using electrodes over the wrists and around the eyes (vertical and horizontal for blinks and saccades), and a forehead ground electrode. Eye movements were also recorded using an EyeLink 1000 infrared eye-tracker (SR Research, ON, Canada). Manual responses were collected using a four-button fibre-optic response box.

MEG data were preprocessed using the OHBA Software Library (OSL; ohba-analysis.github.io) and Fieldtrip (Oostenveld et al., 2011). Continuous MEG data were visually inspected to exclude segments containing severe artefacts (e.g., jumps) and interpolate channels that contained frequent severe artefacts. Elekta’s Maxfilter Signal Space Separation algorithm was then applied, together with head movement compensation based on the HPI coil data. In some participants, the trigger signal used to indicate epochs of interest spread into the MEG channels. Therefore, timepoints in the data corresponding to stimulus triggers were linearly interpolated within a 15ms window around trigger onset; to be consistent, this correction was done in all participants. Data were then band-pass filtered at 0.1-40 Hz, downsampled to 250 Hz, and epoched from -2000ms to 2000ms relative to grating onset. Eye-blink- and cardiac-related variance was removed by performing Independent Component Analysis (ICA) on the MEG data using the FastICA algorithm and removing any components significantly correlated (using α = .05) with vertical EOG and ECG time-courses above a threshold of Pearson’s r = 0.1. Horizontal EOG, EyeLink, and MEG data were then visually inspected to reject trials containing saccades or other muscle artefacts.

### Magnetic Resonance Imaging (MRI) data acquisition

To create subject-specific forward models for MEG source reconstruction, existing structural 3T MRI scans for 16 participants were obtained from previous studies in accordance with data sharing procedures. In addition, new structural MRI scans were acquired for 19 participants on a separate day at the Oxford Centre for Functional MRI of the Brain (FMRIB) using a Siemens Magnetom Prisma 3T scanner, with a T1-weighted MP-RAGE sequence (1×1×1mm isotropic voxels, 256×234×192 grid, echo time = 3.94ms, inversion time = 912ms, recovery time = 1900ms). For this data, participants’ noses were included in the field of view to improve registration with the Polhemus head shape and MEG coordinates. MRI data from two participants were not acquired due to early study exclusion and a scheduling issue, respectively. For the latter participant, an average, age-matched (20-24-year-old) brain template was used instead (Richards et al., 2016).

### Behavioural data analysis

Behavioural data were analyzed in MATLAB and in R using RStudio (RStudio Inc., Boston, MA, USA) with the *ez* package (Lawrence, 2016). Accuracy in the reward learning task (Figure 2a) was quantified as percent of choices of the higher reward option and in the attention task as percent of correct target-discrimination responses. Reaction time (RT) analyses (Figure 2b) were performed only on correct trials and using the median as the subject-level summary statistic. Behavioral data were analyzed using repeated-measures analysis of variance (ANOVA), followed by pairwise comparisons. Tests were corrected for multiple comparisons using the Holm-Bonferroni method where relevant. Learning curves of accuracy and RT in the reward learning task were computed by smoothing each participant’s trial-wise data with a 10-trial moving average window within each session and concatenating sessions. Curves for high reward choices include both High vs. Zero and High vs. Low reward trials.

### Multivariate pattern analysis (decoding)

For decoding analyses in both tasks, the main epoch of interest was the 500ms after reward stimulus onset, which provided a window to analyze early representations of reward-associated stimuli (“options” in the reward learning task, “reward cues” in the attention task) in task-relevant and irrelevant contexts. In the attention task, this epoch in the 500/1000ms SOA conditions enabled us to examine the response to the reward stimuli separately from the response to target processing. For completeness, we also examined reward stimulus processing in the 0-500ms after target onset in the 500/1000ms and 0ms SOA conditions.

Decoding was performed on broadband MEG gradiometer data. The true spatial dimensionality of the MEG data after the Maxfilter preprocessing algorithm is ∼64 rather than the 204 gradiometer channels (Woolrich et al., 2011). Therefore, to reduce the number of redundant features for decoding, we first performed Principal Component Analysis (PCA) on the data to extract a subset of dimensions (70) that explain nearly 100% of the variance. Note that this simply re-expresses the data using a smaller set of features and does not reduce the number of informative dimensions. Using the 70-feature × *N*-trial × *T*-time data matrix as input, we conducted multivariate pattern analysis using Mahalanobis distance with 5-fold cross-validation, an approach closely related to linear discriminant analysis (LDA) (Walther et al., 2016; Wolff et al., 2015, 2017). Specifically, to discriminate between two classes, we split all of the data into five random groups (“folds”) of trials, iteratively using one as a testing set and the remaining as training data (and repeating the splitting procedure 50 times to ensure stable results). We computed Mahalanobis distances between left-out test trials at a given timepoint and the means of *class A* and *B* from the training data at the same timepoint. Mahalanobis distance is given by:

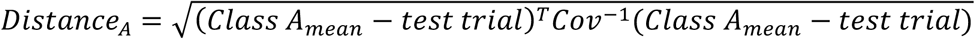

where *Cov*^−1^ is the inverse feature covariance estimated on all training data from both classes (per time-point, across trials) using a shrinkage estimator (Ledoit and Wolf, 2004). The feature covariance captures the correlated noise between sensors or voxels irrespective of condition, and thereby improves decoding reliability (Kriegeskorte et al., 2006; Walther et al., 2016). We then computed the difference between the distances to *Class A* and *Class B*, always subtracting the within-class distance from the between-class distance:

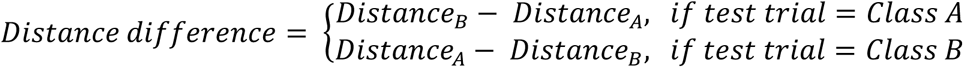

which leads to positive differences with better decodability (Wolff et al., 2017). This was repeated for all trials and timepoints and results were smoothed across time with a Gaussian smoothing kernel (σ = 16ms), to obtain a measure of trial-wise pattern decodability across time. Note that although this distance measure can be binarized and converted to accuracy over trials (as in LDA), we preserved the parametric distance value as a more informative measure of similarity/dissimilarity between train and test patterns (Walther et al., 2016).

We quantified neural pattern separation for various task parameters during the reward learning and attention tasks: reward stimulus location (left versus right; *location decoding*), reward stimulus identity (e.g., high reward stimulus 1 versus high reward stimulus 2; *identity decoding*), and stimulus-independent value (high reward versus low reward; *value decoding*). Location decoding was performed on data collapsed across stimulus identity, and compared between high and low reward trials. Identity decoding was performed separately for each location, with the results averaged over location, and compared between high and low reward trials. For value decoding, we employed a cross-generalisation approach, in which we trained a classifier using one pair of reward stimuli (e.g., high reward 1 vs. low reward 1), and tested performance on another (e.g., high reward 2 vs. low reward 2), collapsing across locations. Value decoding was therefore independent of location and identity, and attributable only to the difference in value between stimuli.

For location and identity decoding in the reward learning task (Fig. 3a-c), the high reward condition included both High vs. Low reward and High vs. Zero reward trials. Value decoding analysis (Fig. 3b) excluded the High vs. Low reward trials to ensure interpretability of results; to obtain equal trial numbers between High vs. Zero and Low vs. Zero trials, we subsampled the latter using an odd/even split. Attention task decoding was performed separately on the 500/1000ms and the 0ms SOA trials.

For statistical testing, we averaged pattern decodability across trials for each participant and tested it at the group level against zero or between conditions using cluster-corrected sign-permutation tests, with a cluster-forming threshold of α = 0.05 and 10,000 permutations. Baseline data (i.e., before stimulus onset) were not included in permutation testing.

To test how location decoding evolved across trials during the reward learning task as a function of reward (Fig. 3b), for each participant, we computed the Spearman rank correlation between trial number and peak location decoding in each trial, separately for high and low reward. We tested the average Fisher z-transformed correlation across participants against zero and between reward conditions using one-sample and paired t-tests, respectively. The time of peak decoding was defined based on the group-level decoding results. Note that due to rejection of trials with artefacts, the number of trials included in this analysis is less than in the learning curves presented in Figure 2.

### Median split analyses

To test the relationship between the observed decoding and the attentional capture effect in behaviour, we split participant decoding results based on the magnitude of their RT reward × congruency interaction effect across all three SOAs. Decoding differences between groups were tested using independent-samples t-statistics and cluster-corrected group-permutation tests, in which observed group differences were compared to a null distribution of differences generated from random group partitions (cluster-forming α = 0.05; 10,000 permutations).

### Time-resolved correlation analyses

To complement the median split analyses, we performed time-resolved Spearman rank correlations between the behavioural attentional capture effect and value modulation in the decoding. Correlations were computed at each timepoint, with statistical significance tested using cluster-corrected permutation tests in which the observed correlation coefficients were compared to a null distribution of coefficients generated from independent shuffles of behavioural and neural participant data (cluster-forming α = 0.05; 10,000 permutations). Additionally, we repeated these analyses after excluding bivariate outliers using a robust multivariate projection method and the box-plot rule (the skipped correlation method in the Robust Correlation Toolbox; Pernet et al., 2012). Outliers were identified and excluded at each timepoint and were also excluded during permutation testing. Post hoc power was calculated using G*Power, assuming a null hypothesis of no correlation (rho = 0), alpha = 0.05, a two-tailed test, and the current sample size of n = 30 (Faul et al., 2009).

### Source reconstruction

Co-registration of MEG sensors and structural MRI was done using OSL (ohba-analysis.github.io). Structural MRI data were bias-corrected and linearly registered to the Montreal Neurological Institute (MNI) space, enabling source reconstruction and analysis to be completed in this common reference space. The MRI scalp surface was extracted and smoothed and fiducial points were manually marked. MRI fiducials were registered to the MEG fiducials using linear transformation and subsequently refined using the iterative closest point algorithm applied to the entire MRI scalp surface and set of Polhemus head shape points. Source reconstruction on MEG gradiometer data was performed in Fieldtrip using a single-shell volume conduction model (based on a mesh of the brain-skull boundary; Nolte, 2003), a leadfield matrix generated from a 10mm-spaced template grid, and linearly constrained minimum variance (LCMV) beamforming (Van Veen et al., 1997). To obtain whole-brain virtual channel data, beamforming filters were applied to time-domain data and, for each gridpoint, the three-direction dipole data were projected to the direction explaining the most variance using singular value decomposition.

### Source-space searchlight decoding

To visualize brain regions involved in decoding, we performed searchlight decoding on whole-brain virtual channel data in source space in 50ms steps, from 250-500ms (location decoding), 0-500ms (identity decoding), and 300-500ms (value decoding) after cue onset. Specific timesteps were chosen because running the full time-course was computationally intensive and difficult to visualise in both the spatial and temporal domain. These timesteps were chosen as they were evenly spaced points corresponding approximately to the significant value modulation of decoding in sensor space, or in the case of identity decoding, to the entirety of the cue epoch. A ∼23mm-radius searchlight sphere was used to define features (grid points) for decoding. The sphere was enlarged at the edges of the brain to maintain an approximately constant number of grid points per sphere (across all spheres, M = 62 grid points, SD = 6 grid points). To reduce extensive computation time, the same feature covariance estimate was used for decoding at each time point, which was computed on baseline data (averaged from -100ms to -50ms relative to cue onset). Additionally, the five-fold split into training and testing data was repeated ten times to obtain stable results. All other aspects of decoding were identical to that described above. For statistical testing, the resultant distance differences were converted into t-statistics and cluster-corrected at the whole-brain level using threshold-free cluster enhancement (TFCE) with 10,000 permutations, as implemented in FSL’s *randomise* tool (Smith and Nichols, 2009; Winkler et al., 2014). For visualization, thresholded t-value maps were first interpolated to a 2mm MNI volume using linear interpolation and then interpolated to the Conte69 inflated surface using trilinear interpolation in Fieldtrip (Marcus et al., 2011; Oostenveld et al., 2011).

### Univariate analysis of event-related fields (ERFs)

To complement the decoding analyses and for comparability with prior research, we conducted univariate analysis of the N2pc and P1 components. Data were baseline-corrected using a 200ms pre-stimulus window. To be optimally sensitive to potential value modulation of evoked components, we first selected sensors for each component using an orthogonal contrast—activity averaged across high and low reward and across all SOA conditions—and then tested the value modulation within those sensors of interest (e.g., Donohue et al., 2016; Luque et al., 2017; MacLean and Giesbrecht, 2015a).

For the N2pc analysis, we identified sensors of interest for each participant, by computing event-related averages for left and right reward cue trials. We then combined planar gradiometers using the root mean square (RMS) approach and computed the difference wave between the cue sides at each sensor. Finally, we identified the five posterior sensors in each hemisphere showing the largest N2pc amplitude averaged within a 200-325ms post-stimulus time window of interest (Itthipuripat et al., 2015). Using these sensors, we compared the strength of the reward-cue-related N2pc between high and low reward conditions for the combined 500/1000ms SOA and the 0ms SOA conditions. For each reward condition, we computed averages separately for left and right reward cue trials; combined planar gradiometers using RMS; averaged across reward cue sides within contralateral and ipsilateral sensors; and computed the sensor-averaged difference between contralateral and ipsilateral conditions (i.e., the N2pc). We then compared the N2pc amplitude, averaged within the 200-325ms window, between high and low reward conditions using a paired t-test.

Identifying sensors for the P1 analysis was similar to that described above, except we additionally allowed for variation of the P1 latency across participants by calculating the difference wave for each sensor within a time window identified specifically for that sensor and participant (MacLean and Giesbrecht, 2015a). To identify the individual P1 latencies, we computed an average across all trials; combined planar gradiometers using RMS; and for each sensor, used the *findpeaks* function from Matlab’s Signal Processing toolbox to identify the latency of the largest peak within a 75-200ms window (see Luque et al., 2017; MacLean and Giesbrecht, 2015a). If no peak was present, we took the latency of the maximum value within that window. The final time window was a 50ms window around the identified peak (Luque et al., 2017; MacLean and Giesbrecht, 2015a). The mean P1 latency (averaged across sensors and participants) was 156ms (SD = 16ms) (similar to Luque et al., 2017). We tested for the value modulation of the P1 in a manner identical to that for the N2pc, except that we compared the P1 difference averaged within a 50ms window around the individual peaks identified earlier.

### Code accessibility

Analysis scripts and data will be made available prior to publication.

## Results

### Reward learning task: Learning stimulus-reward contingencies

We first confirmed that participants learned the stimulus-reward contingencies in the reward learning task. Participants won an average of £44.67 (SD = £2.89) out of a possible £48. Choice accuracy was sensitive to the value difference between options, with a significant main effect of choice type (*high vs. zero*, H-Z; *high vs. low*, H-L; *low vs. zero*, L-Z) on accuracy, F_(2,58)_ = 7.43, p = 0.006. Participants were more accurate on H-Z than L-Z trials, t_(29)_ = 4.36, p < 0.001, Cohen’s d = 0.8 (Fig. 2a). There was no difference between H-Z and H-L, t_(29)_ = 1.37, p = 0.18, or H-L and L-Z trials, t_(29)_ = 2.06, p = 0.1. RT was similarly sensitive to value difference, with a significant main effect of choice type on RT, F_(2,58)_ = 41.4, p < 0.001. Participants were faster on H-Z and H-L choices than L-Z choices, t_(29)_ = 7.69, p < 0.001, Cohen’s d = 1.4 and t_(29)_ = 6.07, p < 0.001, Cohen’s d = 1.1, respectively (Fig. 2b). There was no difference between H-Z and H-L choices, t_(29)_ = 1.14, p = 0.26.

Previous work has dissociated the effects of irrelevant reward history and reward-independent selection history on attentional capture (MacLean and Giesbrecht, 2015b). The latter phenomenon reflects findings showing that merely selectively attending to a stimulus can affect subsequent attention to it, irrespective of reward associations (Awh et al., 2012; MacLean and Giesbrecht, 2015b). To control for selection history, we doubled the number of Low vs. Zero reward trials to match the frequency of choosing low reward options with that of high reward options. We confirmed that choice frequency (i.e., how often participants selected a given reward option) did not differ between high and low reward options (note that this is correlated with but not identical to trial-type frequency, which was fixed across participants). There was a main effect of stimulus value on choice frequency, F_(2,58)_ = 721.05, p < 0.001. Participants chose the high and low reward options more frequently than the zero reward stimuli (High vs. Zero, t_(29)_ = 35, p < 0.001, Cohen’s d = 6.39; Low vs. Zero, t_(29)_ = 28.26, p < 0.001, Cohen’s d = 5.15). There was no difference between high and low reward options, t_(29)_ = 0.6, p = 0.55 (Fig. 2c), demonstrating the absence of a choice frequency (i.e., selection history) effect.

### Reward learning task: reward stimulus processing in a task-relevant context

Next, we tested the feasibility of time-resolved decoding of reward stimulus representations in the reward learning task, a context in which these stimuli (i.e., *options* for decision-making) are relevant and rewarded. These analyses focused on the fixation epoch, in which participants saw the options but were not yet able to respond.

We applied multivariate pattern analysis to discriminate between trials in which the higher valued option appeared on the left versus the right of the fixation cross, separately for the high and low reward trials (*location decoding;* Fig. 3a). Given the obvious perceptual differences between left and right option trials, we expected the decoding analysis to reflect this. Crucially, however, these perceptual differences should be identical in high and low reward trials, and therefore any differences in decoding between these conditions are attributable solely to the reward associated with each type of option. As expected, for both high- and low-reward options, we could decode option location from ∼104ms and ∼136ms onwards, respectively (each p < 0.001, cluster-corrected permutation test), in line with the clear perceptual difference between left and right cue presentation trials. Critically, the decoding was significantly stronger for the high-than the low-reward options from ∼184ms onwards (p < 0.001), indicating that learned value modulated location decodability in this task-relevant context (Fig. 3a). This is consistent with an effect of reward on spatial attention (Anderson, 2016; MacLean et al., 2016; Moore and Zirnsak, 2017).

Given that participants gradually learned to locate and select the higher reward stimulus, we examined how location decoding emerged across trials as a function of associated reward. We extracted trial-wise distance differences from the peak time point of maximal decoding in the trial-averaged data in the high and low reward conditions (i.e., Fig. 3a) and tested for a relationship with trial number. In line with participants’ learning, location decoding significantly increased over trials in the high reward condition (t-test on the participant-wise, Fisher z-transformed Spearman rank correlations; t_(29)_ = 8.68, p < 0.001; see also Fig. 3b top) but not in the low reward condition (t_(29)_ = 1.95, p = 0.06; see also Fig. 3b bottom). Correlations were stronger in the high reward compared to the low reward condition (t_(29)_ = 5.7, p < 0.001; Fig. 3b inset). Thus, the emergence of location decoding across trials also differed as a function of associated reward, consistent with the idea that reward learning gradually shapes selective attention (Anderson, 2016; Chelazzi et al., 2013).

Next, we applied a similar multivariate pattern analysis to discriminate between the two stimulus identities within each reward condition (e.g., high reward 1 vs. high reward 2; *identity decoding*; Fig. 3c). Identity could be decoded from ∼80ms and ∼104ms onwards for high and low reward stimuli, respectively (p < 0.001, cluster-corrected permutation test), but with no difference between reward conditions (p = 0.417), suggesting that learned value did not modulate the sensory representations of these stimuli in this task (Fig. 3c).

Lastly, we sought to decode a reward value signal, independent of location or identity. We used a cross-generalisation approach, in which we trained a classifier using one pair of reward stimuli (e.g., high 1 vs. low 1), and tested performance on another (e.g., high 2 vs. low 2), collapsing across locations (e.g., Kahnt et al., 2010). Any significant decoding that emerges is therefore independent of location and identity, and so attributable to the difference in value between stimuli (*value decoding;* Fig. 3d). We found a significant value signal from ∼260ms onwards (p = 0.001, cluster-corrected permutation test), representing the available reward independently of stimulus location or identity (Fig. 3d).

### Attention task: Attentional capture by stimuli associated with reward history

Next, we probed how stimulus-reward associations learned in the reward learning task captured attention in the attention task and whether attentional capture was modulated by reward value and/or by the SOA. We ran a repeated-measures ANOVA on RT with three factors: *reward* (high, low), *congruency* (congruent, incongruent), and *SOA* (0, 500, 1000ms). There was a main effect of congruency, F_(1,29)_ = 12.46, p = 0.001 and a congruency × reward interaction, F_(1,29)_ = 8.9, p = .006. Participants were significantly slower in the high reward incongruent relative to the congruent condition, t_(29)_ = 3.77, p = 0.002, Cohen’s d = 0.69, with no such difference between the low reward incongruent and congruent conditions, t_(29)_ = 1.77, p = 0.087, Cohen’s d = 0.32. Importantly, although there was a main effect of SOA, F_(2,58)_ = 44.33, p < 0.001 (indicative of a general preparatory effect), there were no interactions between SOA and reward or congruency, all F_(2,58)_ < 1, p > 0.3. Thus, VDAC effects in reaction time were consistent across SOAs. For completeness, we tested for reward × congruency effects in each SOA. Across SOA conditions, participants were slower in the high-reward incongruent trials than the congruent trials: *0ms SOA*, t_(29)_ = 4.55, p = 0.001, Cohen’s d = 0.83; *500ms SOA*, t_(29)_ = 3.15, p = 0.019, Cohen’s d = 0.57; *1000ms SOA*, t_(29)_ = 2.8, p = 0.036, Cohen’s d = 0.51 (Figure 4a). There was no difference between low-reward incongruent and congruent trials, all t_(29)_ < 2, p > 0.2 (Figure 4a).

A congruency effect on reaction time could be driven both by faster responses in congruent trials, and/or slower responses in incongruent trials, compared to baseline. Figure 4a suggests that both might be the case, and we next tested this explicitly. Given the absence of interactions with SOA, we collapsed across this condition and performed one-sample t-tests for each congruency and reward condition. In high-reward trials, participants showed both a congruent (t_(29)_ = 3.14, p = 0.004, Cohen’s d = 0.57) and incongruent effect (t_(29)_ = 2.59, p = 0.015, Cohen’s d = 0.47), suggesting that they were respectively sped up or slowed down by the presence of a congruent or incongruent high-reward stimulus. These effects were not present in the low-reward conditions (t_(29)_ = 1.2, p = 0.238; and t_(29)_ = 1.38, p = 0.178, respectively).

We then tested whether the effect of congruency on RT decayed over trials, as participants repeatedly encountered the unrewarded and irrelevant stimuli. This would reflect an extinction of the stimulus-reward association as reward becomes unavailable, as widely observed in learning paradigms (Todd et al., 2014). Focusing on the high reward condition and collapsing across SOAs, we split trial data into two halves per session and computed a difference between congruent and incongruent RTs as an aggregate measure of attentional capture for each half. A *session* × *half* repeated-measures ANOVA showed no significant main effects or interactions, all F_(1,29)_ < 2.8, p > 0.11, suggesting that the effect did not reliably extinguish across trials within the timeframe of this experiment (see also Figure 4b).

We also analysed accuracy difference scores. There was a significant main effect of congruency, F_(1,29)_ = 9.25, p = 0.005; a main effect of SOA, F_(2,58)_ = 3.17, p = 0.049; a reward × congruency interaction, F_(1,29)_ = 20.11, p = 0.0001; and a reward × congruency × SOA interaction, F_(2,58)_ = 3.21, p = 0.047. In the high reward 500ms and 1000ms SOA conditions only, participants were less accurate in the incongruent than in the congruent trials, t_(29)_ = 3.75, p = 0.005, Cohen’s d = 0.68 and t_(29)_ = 3.45, p = 0.009, Cohen’s d = 0.63. There were no differences in the 0ms SOA or in any low reward condition, all t_(29)_ < 2, p > 0.9. Thus, attentional capture also influences accuracy, at least in the longer SOA conditions, with no evidence of a speed-accuracy trade-off.

### Reward cue location is decodable in a task-irrelevant context and modulated by value

Next, we probed the neural correlates of the attentional capture effect we observed in behaviour. We aimed to identify neural signatures of task-irrelevant reward history, independently of target processing. Similar to the analysis of the reward learning task data, we performed time-resolved decoding of the location and identity of the reward cues, as well as a stimulus-independent value signal. This enabled us to probe the time-course of any effects of reward history, such as modulations in early visual processing. Our primary focus was the cue epoch, prior to the onset of task-relevant gratings, as this period allowed for the analysis of task-irrelevant reward cue processing without the interference of target processing. We specifically focused on the first 500ms after reward cue onset as this allowed us to collapse across the 500ms and 1000ms SOA trials.

We first looked at location decoding as the most direct correlate of the spatially-specific attentional capture which would subsequently impact target processing and related behaviour. As before, we found significant decoding of the cue location for both high (p < 0.001, cluster-corrected permutation test) and low reward cues (p < 0.001; Fig. 5a). Critically, this decoding was modulated by reward value, with stronger decoding for high compared to low reward cues starting from ∼260ms (p < 0.001; Fig. 5a). This effect is solely attributable to the reward history that these stimuli acquired during the reward learning task; in the current context, these stimuli are unrewarded and irrelevant.

**Figure 5.**
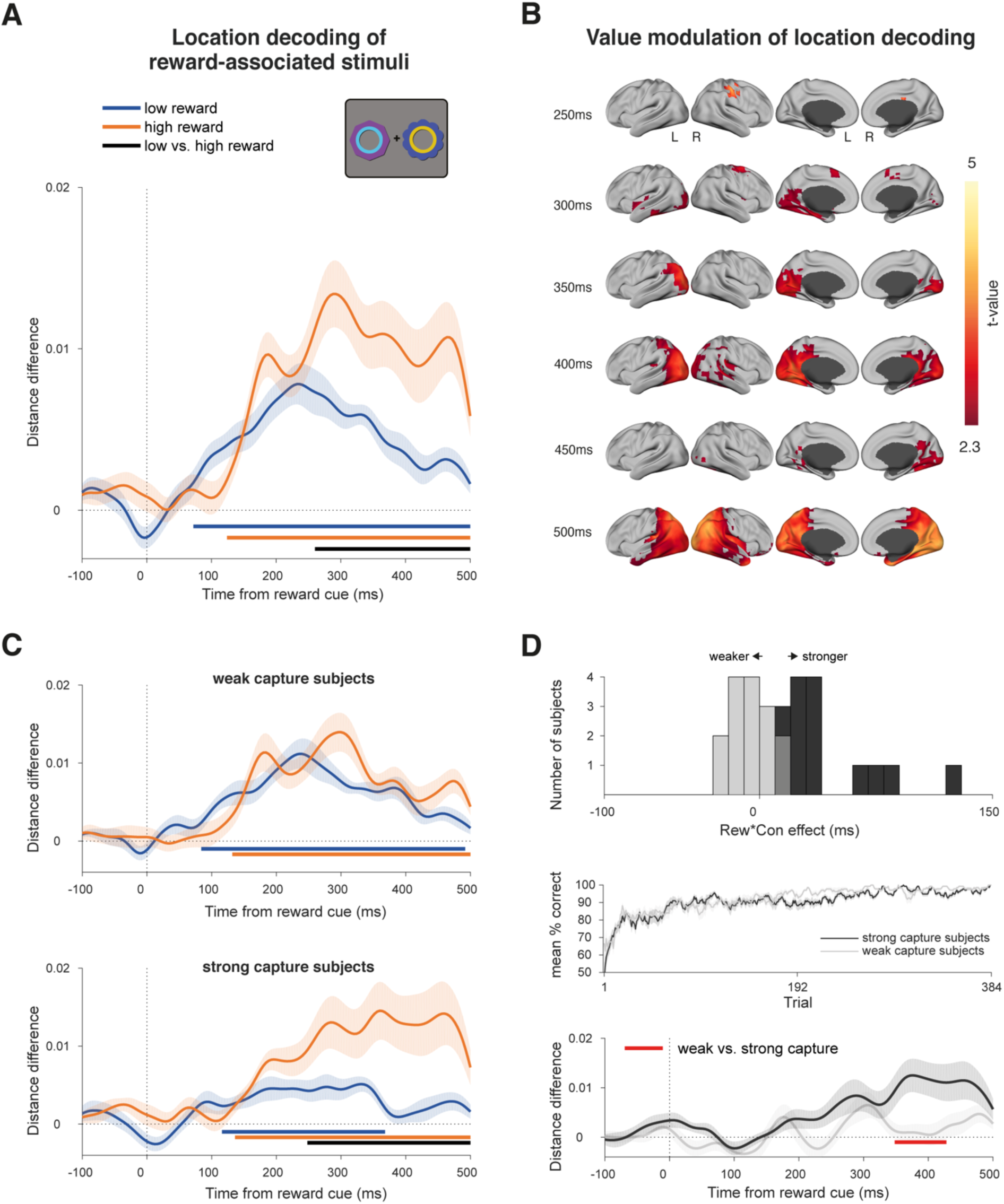
Decoding reward cue location in the attention task during the cue epoch of the 500ms and 1000ms SOA conditions. **(a)** Decoding of reward cue location during the cue epoch for high- and low-reward stimulus trials. Curves depict decoding averaged across trials and participants for high and low reward options. Bars indicate significant decoding (p < 0.05, cluster-based permutation test across time) for each reward condition (orange, blue bars) and between reward conditions (black bar). Shaded regions indicate SEM. **(b)** Significant value modulation of location decoding in a source-space searchlight analysis. Maps depict t-statistics at selected time-points during the cue epoch, and are thresholded at p < 0.05, whole-brain cluster-corrected using TFCE. **(c)** Same decoding as (a) but performing a median split on the participant-level data according to each participant’s reward × congruency behavioural effect (i.e., value-modulated attentional capture effect; see Figure 5d). *Top*: the “weak capture” group. *Bottom*: the “strong capture” group. **(d)** Behavioral and decoding comparison of each participant group from (c). *Top*: histogram of participants’ reward × congruency behavioural effect (weak capture in grey, strong capture in black). *Middle*: learning curves for each group. Same color conventions as above. *Bottom*: Differences between high- and low-reward location decoding (orange minus blue lines) for each group from (c). Same color conventions as above. Red bar indicates significant difference between groups (p < 0.05, independent-samples cluster-based permutation test across time).

We sought to determine the anatomical source of this value modulation. We projected the sensor-space data onto a source-space grid using a linearly constrained minimum variance (LCMV) beamformer, and ran the same location decoding analysis within a searchlight sphere across the brain, in 50ms bins from 250-500ms after cue onset. Value modulation began around the right posterior parietal cortex, followed by the inferior temporal and visual cortex and more widespread modulation in posterior parietal cortex, and along the temporal lobe (p < 0.05, whole-brain cluster-corrected using TFCE; Fig. 5b).

We then tested whether this value modulation was related to the value modulation of the behavioural attentional capture effect. We conducted a median split of the participants based on the extent to which their attentional capture was modulated by reward value (i.e., the strength of the reward × congruency interaction effect on RT; Fig. 5d top panel). Strong capture participants (i.e., with stronger value modulation of the attentional capture effect) showed accompanying value modulation of the location decoding during the cue epoch starting from ∼244ms (p < 0.001, cluster-corrected permutation test; Fig. 5c bottom panel), while weak capture participants did not (p = 0.206; Fig. 5c top panel), with a significant difference between these two groups (p = 0.025, independent-samples, cluster-corrected permutation test; Fig. 5d bottom panel). A time-resolved Spearman rank correlation analysis between the behavioural VDAC effect and the neural value modulation across participants mirrored these results (cluster from 348-408ms, average within-cluster rho = 0.504, p = 0.031, cluster-corrected permutation test). We also confirmed that these results were not driven by outlying participants using a skipped rank correlation which formally excludes bivariate outliers (cluster from 352-416ms, average within-cluster rho = 0.508, p = 0.004; for details, see methods and Pernet et al., 2012). Finally, we note that according to post-hoc power calculations, with the current sample size of n = 30 and alpha = 0.05, there is 80% power to detect a correlation of rho = 0.5.

We defined the above strong and weak capture groups based on the extent to which reward value modulates the behavioural effects of attentional capture, but they might also differ in how they learn reward associations (e.g., Jahfari and Theeuwes, 2017). We therefore compared their learning curves from the reward learning task, but did not find any significant differences (p = 0.410, independent-samples, cluster-corrected permutation test across trials; Fig. 5d middle panel).

Next, we tested whether the cue location decoding persisted into the target epoch, which would suggest that participants maintain traces of the irrelevant reward cue location even while discriminating the oriented gratings. We repeated the location decoding on 500ms and 1000ms SOA trial data, now time-locked to the grating onset. We again found significant decoding of cue location for both high (p < 0.001, cluster-corrected permutation test) and low reward cues (p < 0.001), which was also modulated by reward value (p < 0.001; Fig. 6a). Note that the significant decoding prior to grating onset (the typical “baseline” period) occurs because this period overlaps with the preceding cue epoch. As above, we tested whether this value modulation during the target epoch related to the behavioural attentional capture effect. A median split analysis (described above) showed no significant difference in value modulation of location decoding between strong and weak capture participants (p = 0.283, independent-samples, cluster-corrected permutation test). Strong capture participants showed significant value modulation (p = 0.002, cluster-corrected permutation test), as did weak capture participants (p = 0.049; Fig. 6b). A complementary time-resolved correlation analysis (as above) also showed no significant effects during this time period (Spearman rank correlation: p = 0.336, cluster-corrected permutation test; skipped rank correlation: no data exceeded the cluster-forming threshold).

**Figure 6.**
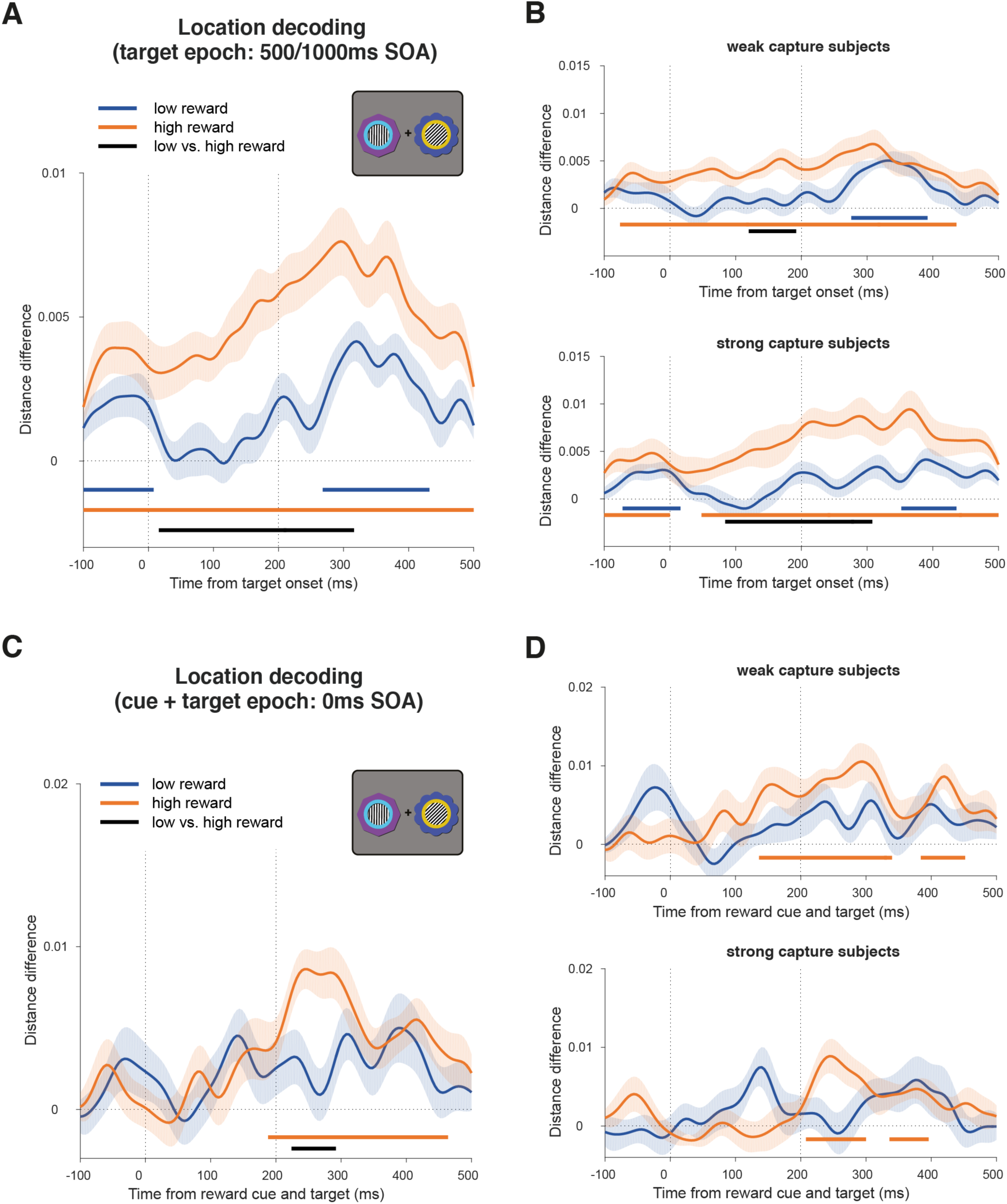
Decoding reward cue location in the attention task during the target epoch. **(a)** Decoding of reward cue location in the target epoch in the 500ms and 1000ms SOA trials. Note that significant decoding in the baseline period is due to the temporal overlap with the cue epoch. Curves depict decoding averaged across trials and participants for high and low reward options. Bars indicate significant decoding (p < 0.05, cluster-based permutation test across time) for each reward condition (orange, blue bars) and between reward conditions (black bar). Shaded regions indicate SEM. Vertical dashed lines respectively indicate onset of gratings and offset of gratings and reward cues. **(b)** Same decoding as (a) but performing a median split on the participant-level data according to each participant’s reward × congruency behavioural effect (see Figure 5c-d for details). **(c-d)** Same analysis as (a-b) but for the 0ms SOA condition (simultaneous cue and target onset). Vertical dashed lines respectively indicate onset and offset of gratings and reward cues.

We also performed the same location decoding analysis in the 0ms SOA condition, in which reward cues and gratings are presented simultaneously. We found significant decoding of the cue location for high (p < 0.001, cluster-corrected permutation test) but not low reward cues (p = 0.121), and a significant value modulation from ∼224ms (p = 0.024; Fig. 6c). As before, we conducted a median split analysis (described above). However, we did not find value modulation of the location decoding when analysing each group separately (strong capture participants: p = 0.123; weak capture participants: p = 0.519; Fig. 6d). A complementary time-resolved correlation analysis (as above) also showed no significant effects (Spearman rank correlation: p = 0.175; skipped rank correlation: p = 0.427). Note that these analyses should be interpreted with caution because of simultaneous cue and target processing in this condition as well as the reduced within-subject power, with half as many trials here as in the combined 500ms and 1000ms SOA decoding.

We have used here linear multivariate methods as a sensor- and amplitude- agnostic method of analysing cue-related activity. To ensure that the above results are not specific to our method, and for comparability with prior research, we also conducted univariate analysis of the event-related field (ERF) responses. Based on prior research on VDAC, we focused on value modulation of the early sensory-evoked P1 and the later N2pc components (Hickey et al., 2010; Itthipuripat et al., 2015; Luque et al., 2017; MacLean and Giesbrecht, 2015a; Qi et al., 2013). To be optimally sensitive to value effects, we performed these analyses within time windows of interest based on prior research, and in sensors identified for each participant using an orthogonal contrast (for details, see Materials and Methods; Donohue et al., 2016; Itthipuripat et al., 2015; Luque et al., 2017; MacLean and Giesbrecht, 2015a). Consistent with the location decoding results, we found a value modulation of the N2pc in the 500ms and 1000ms SOA condition (high vs. low reward N2pc amplitude: t_(29)_ = 2.186, p = 0.037; Figure 7a) and in the 0ms SOA condition (high vs. low reward N2pc amplitude: t_(29)_ = 2.362, p = 0.025; Figure 7b). In contrast, we did not observe any value modulation of the P1 component in the 500ms and 1000ms SOA condition (high vs. low reward P1 lateralisation: t_(29)_ = 0.621, p = 0.54; Figure 7c), nor in the 0ms SOA condition (high vs. low reward P1 lateralisation: t_(29)_ = 0.176, p = 0.861; Figure 7d). Thus, the timing of value modulation effects is consistent across multivariate and univariate analyses, with an absence of value modulation in early time windows (i.e., the evoked P1 component), but the presence of value modulation in later time windows (i.e., the N2pc component).

**Figure 7.**
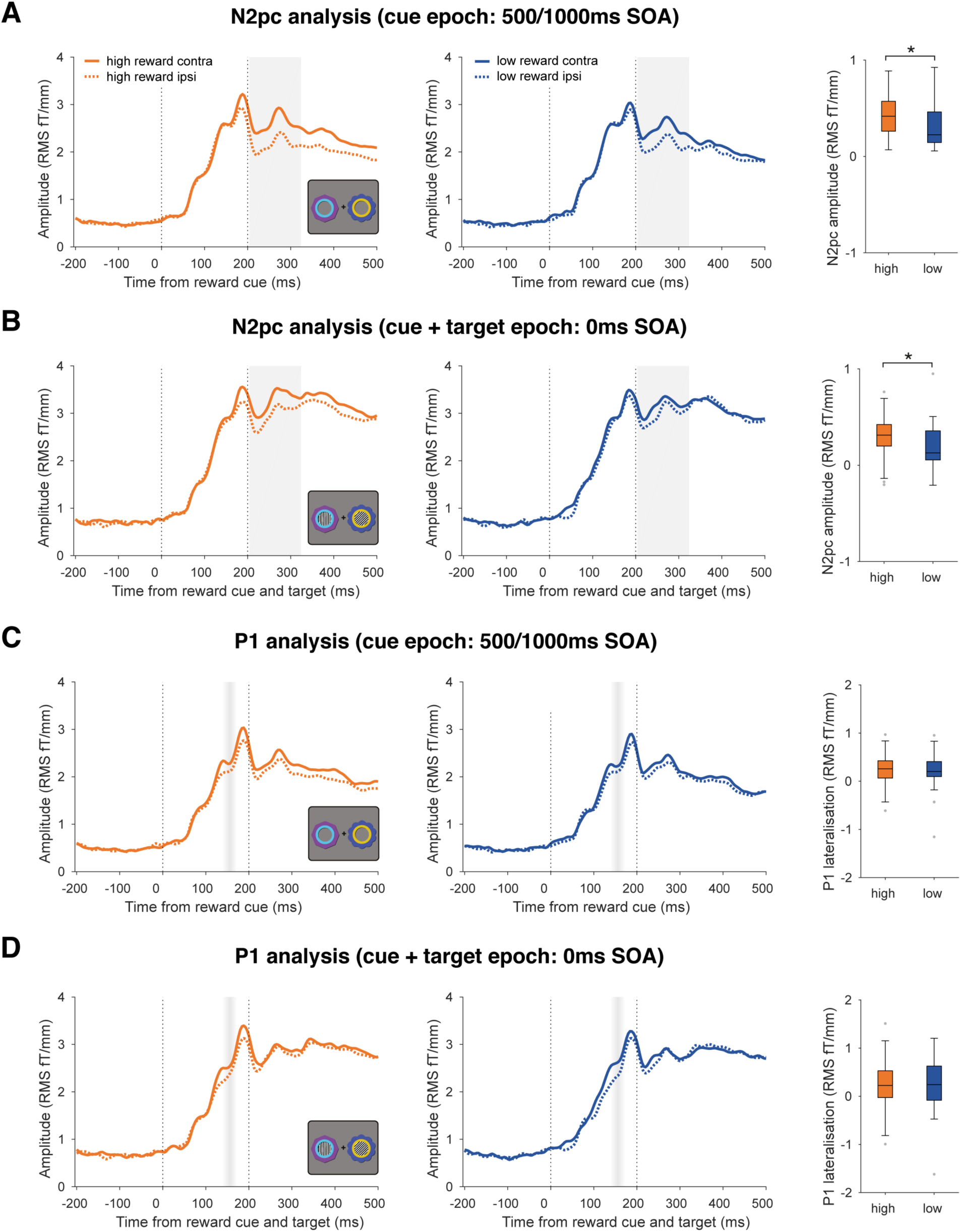
Value modulation of evoked responses. (a) Value modulation of the N2pc in the 500/1000ms SOA trials. *Left and middle panels:* Event-related fields (ERFs) from combined planar gradiometer sensors contralateral and ipsilateral to the reward cues, for high and low reward trials (orange and blue lines) in the 500/1000ms SOA condition. The difference between contralateral and ipsilateral lines is the N2pc. Grey shading indicates the 200-325ms time window of interest for the analysis. Vertical dashed lines indicate reward cue onset and offset. *Right panel:* N2pc amplitude (the difference between contralateral and ipsilateral ERFs) for high and low reward conditions, averaged within the time window of interest indicated in grey. Asterisk indicates p < 0.05. Box plots follow same conventions as Figure 2. (b) Value modulation of the N2pc in the 0ms SOA trials. Vertical dashed lines indicate target grating onset and target grating/cue offset. All other conventions identical to (a). (c) No evidence of value modulation of the P1 in the 500/1000ms SOA trials. *Left and middle panels:* ERFs from combined planar gradiometer sensors contralateral and ipsilateral to the reward cues, for high and low reward trials (orange and blue lines) in the 500/1000ms SOA condition. Faded grey shading indicates the mean ± 1 standard deviation of the P1 peak latency across participants (averaged across sensors). Vertical dashed lines indicate reward cue onset and offset. *Right panel:* P1 lateralisation (the difference between contralateral and ipsilateral ERFs) for high and low reward conditions, averaged within a 50ms window around a sensor- and subject-specific P1 peak. Box plots follow same conventions as Figure 2. (d) No evidence of value modulation of the P1 in the 0ms SOA trials. Vertical dashed lines indicate target grating onset and target grating/cue offset. All other conventions identical to (c).

### No evidence that reward cue identity decoding is modulated by value in a task-irrelevant context

In addition to the location decoding of the reward cues (i.e., reflecting spatial selection), it is possible that decoding of the cue identities (putatively reflecting visual feature representations) would be modulated by learned value (Anderson, 2016; Failing and Theeuwes, 2018). Although we did not observe an effect of reward value on identity decoding during the reward learning task, we nevertheless explored this possibility during the attention task. Although we could reliably decode identity for each type of reward cue (high reward, from ∼128ms, p < 0.001; low reward, from ∼124ms, p < 0.001), this was not modulated by value (p = 0.115; Figure 8a). When we performed a median split analysis by behaviour (described above), the strong capture participants did not show any significant value modulation (p = .065; Figure 8b bottom panel). Weak capture participants also did not show any value modulation (p = 0.353; Figure 8b top panel). A time-resolved correlation analysis (as above) also showed no significant effects (Spearman rank correlation: p = 0.228; skipped rank correlation: p = 0.334). A similar pattern was observed when decoding in the 0ms SOA trials, with significant decoding of stimulus identity in high and low reward trials (for both, p < 0.001), but no significant value modulation (no data exceeded the cluster-forming threshold; Figure 8c). Likewise, neither strong nor weak capture participants showed significant value modulation (for both, no data exceeded the cluster-forming threshold candidates; Figure 8d). A time-resolved correlation analysis (as above) also showed no significant effects during this time period (Spearman rank correlation: p = 0.241; skipped rank correlation: p = 0.158).

**Figure 8.**
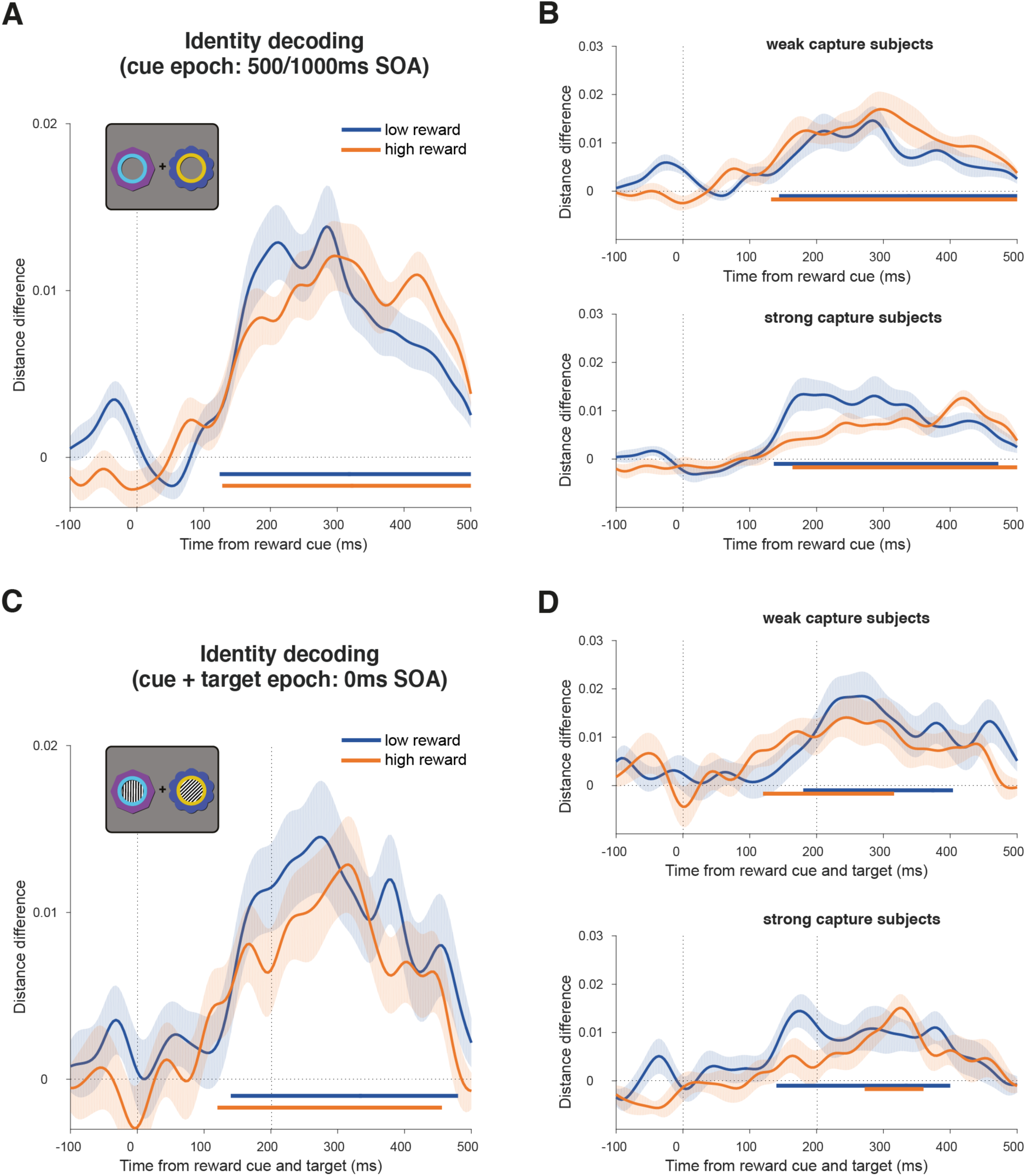
Decoding reward cue identity in the attention task. **(a)** Decoding of reward cue identity during the cue epoch in the collapsed 500ms and 1000ms SOA conditions. Curves depict decoding averaged across trials and participants for high and low reward options. Bars indicate significant decoding (p < 0.05, cluster-based permutation test across time) for each reward condition (orange, blue bars). Shaded regions indicate SEM. Vertical dashed line indicates onset of reward cues. **(b)** Same decoding as (a) but performing a median split on the participant-level data according to each participant’s reward × congruency behavioural effect (see Figure 5c-d for details). **(c-d)** Same analysis as (a-b) but for the 0ms SOA condition. Vertical dashed lines respectively indicate onset and offset of gratings and reward cues.

It is possible that any effect of reward history on identity decoding is confined to a specific anatomical region (e.g., within the ventral visual stream), which our whole-brain sensor-space analysis may have missed. To test this, we repeated the analysis using source-space searchlight decoding across the brain in 50ms time steps from cue onset and tested for significant clusters. This approach enabled a more fine-grained sub-selection of potentially informative signals (i.e., within a searchlight region-of-interest), relative to the whole-brain sensor-space decoding analysis. There were no significant differences between high and low reward trials anywhere in the brain (500ms and 1000ms SOA: p > 0.1; 0ms SOA: p > 0.209; whole-brain cluster-corrected using TFCE). Thus, we did not find any evidence that reward history modulates stimulus identity representations in either a task-relevant, or a task-irrelevant context in the current experiment. Moreover, any hint of an effect is in the opposite direction to that expected, with slightly worse decoding in the high reward condition.

### Decoding stimulus-independent reward value in a task-irrelevant context

Finally, we sought to decode a location- and identity-independent value signal during the attention task, which would provide additional evidence that value information continues to be represented in a task-irrelevant context. Using the same cross-generalisation approach as before, we found significant value decoding in the cue epoch, emerging from ∼336ms (p = 0.003, cluster-corrected permutation test; Fig. 9a). In a median split analysis based on participants’ behaviour (described above), neither the strong nor the weak capture group showed significant decoding by itself (strong capture: p = 0.086; weak capture: p = 0.143), although the strong capture participants appeared to be driving the overall effect (Fig. 9b). The difference between groups was not significant (no data exceeded the cluster-forming threshold, independent-samples, cluster-corrected permutation test). The absence of significant decoding in either of the two subgroups, together with significant decoding across the entire sample, is most likely explained by a lack of statistical power in the former analysis, rather than a true null effect. A time-resolved correlation analysis (as above) also showed no significant effects during this time period (Spearman rank correlation: p = 0.361, cluster-corrected permutation test; skipped rank correlation: p = 0.480). Source-space searchlight decoding of this value signal across the brain showed significant clusters in bilateral visual, left inferior temporal, and left inferior frontal cortex (p < 0.05, whole-brain cluster-corrected using TFCE; Fig. 9c). In the 0ms SOA condition, we found no significant value decoding at any timepoint (p = 0.210, cluster-corrected permutation test).

**Figure 9.**
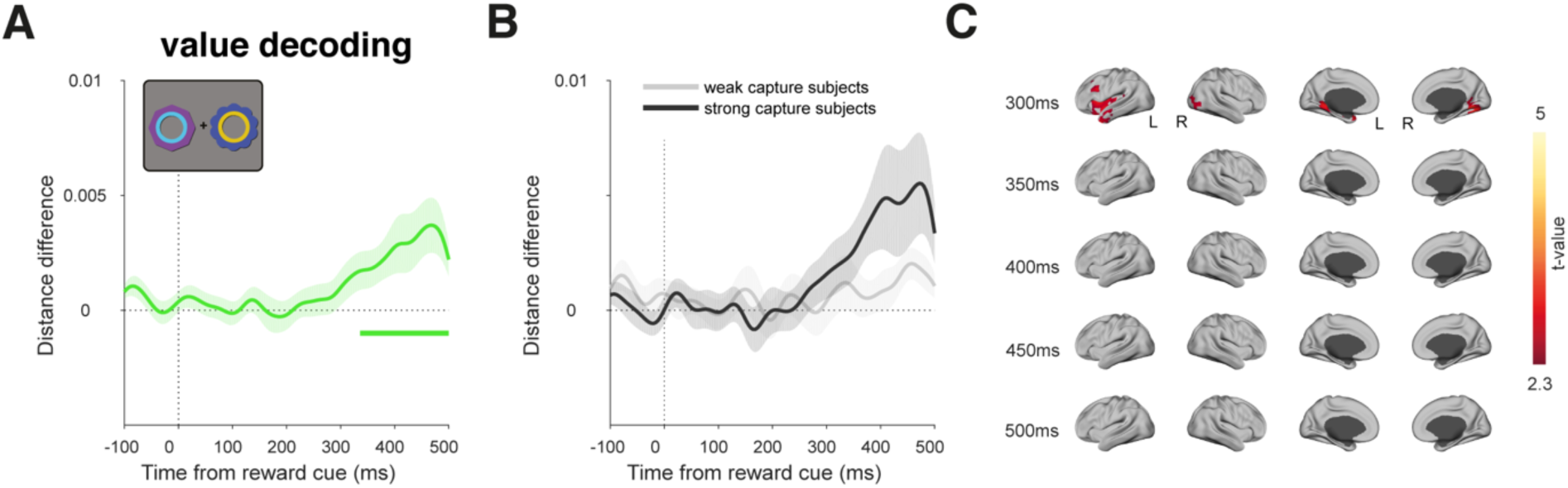
Value decoding in the attention task during the cue epoch of the 500ms and 1000ms SOA conditions. **(a)** Value decoding during the cue epoch in the collapsed 500ms and 1000ms SOA conditions. Curve depicts decoding averaged across trials and participants. Bar indicates significant decoding (p < 0.05, cluster-based permutation test across time). Shaded regions indicate SEM. **(b)** Same decoding as (a) but performing a median split on the participant-level data according to each participant’s reward × congruency behavioural effect (see Figure 5c-d for details). **(c)** Significant value decoding in a source-space searchlight analysis at selected time-points. Same conventions as Figure 5b.

## Discussion

We recorded brain activity using MEG as participants completed a reward learning task to establish stimulus-reward associations, followed by an attention task to measure VDAC. First, we showed a spatially specific behavioural VDAC effect, with reward-associated stimuli respectively speeding or slowing RT in congruent and incongruent trials, consistent with attentional costs and benefits (Failing and Theeuwes, 2014; Posner, 1980). This effect was modulated by reward value, suggesting that it is the associated *value* that is driving capture, rather than a reward-independent selection history (MacLean and Giesbrecht, 2015b). VDAC was robust even with a delay between reward-associated stimuli and targets (see also Failing and Theeuwes, 2015), allowing us to isolate the processing of reward-associated stimuli. Using MVPA, we found that location decoding of task-irrelevant reward cues was modulated by value from ∼260ms after stimulus onset. This neural effect was related to behaviour: participants who had strong VDAC also had larger value modulation of location decoding. Further, we found that this value modulation extended into the target epoch, consistent with the behavioural effects of VDAC observed in the long SOA conditions (500ms and 1000ms). We found an additional value signal in the long SOA conditions, which was independent of the location or identity of the reward stimuli. In contrast, although we could decode the identity of reward cues, this decoding was not modulated by value. This was the case regardless of SOA condition; whether we focused our analysis on participants showing a strong VDAC effect; or whether we performed decoding on whole-brain data or with searchlight analysis.

### Value signals and spatial attention

The timing of the value modulation of location decoding is consistent with a lateralised N2pc signal for reward-associated stimuli, and indeed we found a value modulation of the N2pc in a univariate analysis (Hickey et al., 2010; Itthipuripat et al., 2015; Qi et al., 2013). This suggests an effect of value on spatial attention (Chelazzi et al., 2014; MacLean et al., 2016; Moore and Zirnsak, 2017; Woodman and Luck, 1999). Moreover, localisation of this effect to the posterior parietal cortex (PPC) is in line with theories proposing that regions in this area (e.g., the lateral intraparietal area [LIP] in monkeys) integrate diverse signals, including task information and physical salience, to compute the overall attentional priority of stimuli (Bisley and Goldberg, 2010; Zelinsky and Bisley, 2015). Single-unit recording and fMRI studies suggest that learned reward value is similarly integrated in the PPC (Anderson et al., 2014; Barbaro et al., 2017; Ghazizadeh et al., 2018; Gottlieb and Snyder, 2010). For example, LIP neurons encode learned value despite its interference with goal-oriented behaviour, and such biases transfer to a task-irrelevant context after extensive training (Peck et al., 2009). Interestingly, after the onset of task-irrelevant reward cues, LIP neurons show sustained increases in firing rate for at least 800ms (Peck et al., 2009). This dovetails with our findings that value modulation of location decoding extends into the target epoch and that behavioural VDAC effects are also present at long SOAs. The role of learned value in these effects is demonstrated by the relatively weaker location decoding of low reward cues in both the cue and target epochs, a pattern which mirrors the behavioural results.

The spatial resolution of source localization in the current study was insufficient to confidently distinguish parietal sub-regions (Corbetta and Shulman, 2002). Our findings are also consistent with studies showing that ventral parts of the PPC (e.g., inferior parietal lobule) are involved in the detection of salient events and attentional orienting (Corbetta and Shulman, 2002; Husain and Nachev, 2007). The processing of reward history in this and other parietal sub-regions remains an important open question (Frank and Sabatinelli, 2012).

### Value signals, VDAC, and the stimulus-onset asynchrony

Although we observed a relationship between the VDAC RT effect and the value modulation of location decoding in the 500/1000ms SOA condition, this was not apparent in the 0ms SOA condition. We suggest that this discrepancy stems from practical differences between conditions. Namely, the 0ms SOA condition involves simultaneous processing of reward cue and target stimuli, which may hamper decoding, and is the primary reason that we included an SOA in the paradigm. Moreover, the 0ms SOA condition has half as many trials as the combined 500/1000ms SOA condition, decreasing within-subject power for decoding. Consistent with this interpretation, we did not observe that the VDAC RT effect differed between SOA conditions (in line with Failing and Theeuwes, 2015).

More broadly, we note that cross-subject, brain-behaviour correlations need to be interpreted with caution (Yarkoni, 2009). Nevertheless, our finding of a link between value modulation of stimulus-driven activity and VDAC effects in RT is consistent with previous studies reporting similar effects using fMRI (Hickey and Peelen, 2015) and EEG (Hickey et al., 2010; Qi et al., 2013).

### The latency of value signals and VDAC

The latency of observed value signals is too late to suggest a causal role in modulating visual processing. Attentional modulations arising from frontal and parietal cortex—the putative effects of VDAC—have been shown to already occur by ∼250ms (Anderson, 2016; Buffalo et al., 2010; Foxe and Simpson, 2002). Indeed, robust behavioural effects are also observed when targets are presented simultaneously to the reward stimuli (e.g., 0ms SOA condition; see also Anderson et al., 2011). In these trials, it is likely that any value signal would have to occur earlier than ∼200ms to causally influence visual processing (Buffalo et al., 2010; Foxe and Simpson, 2002). It is possible that there is an early cortical effect which we cannot detect due to a lack of power, or due to the relative insensitivity of MEG to radially-oriented dipoles compared to EEG (Ahlfors et al., 2010). As described earlier, some EEG studies do report value modulation earlier than ∼200ms (Hickey et al., 2010; Luque et al., 2017; MacLean and Giesbrecht, 2015a).

Differences in task design may explain the discrepancies between studies. Hickey et al. (2010) observed a P1 effect using an inter-trial priming paradigm in which the reward history and task relevance of stimuli changed across trials (see also Itthipuripat et al., 2015). Notably, MacLean and Giesbrecht (2015a) did find a P1 effect days after reward learning, but it only occurred in trials in which the reward-associated stimulus directed attention towards the target, suggesting an interaction with task relevance. Similarly, the P1 effect in Luque et al. (2017) occurred in the outcome phase of a reinforcement learning task in which stimuli are task-relevant. In contrast, in the current study and in Qi et al. (2013), who also did not find a P1 effect, the reward-associated stimuli were entirely irrelevant in the attention task. Therefore, it is possible that early cortical value signals may only be present, or at least detectable, when they coincide with current or recent task-related attentional modulation (Failing and Theeuwes, 2018).

### Implications for the visual cortical plasticity account

The visual cortical plasticity account of VDAC posits that value may be directly encoded in cortex via plasticity of stimulus representations in the ventral visual stream (Anderson, 2016; Failing and Theeuwes, 2018; van Koningsbruggen et al., 2016). High-level object, perceptual, and category learning are known to induce plasticity throughout the ventral visual stream and higher-order areas (LeMessurier and Feldman, 2018; Miyashita, 1988; Op de Beeck and Baker, 2010; Schoups et al., 2001; Srihasam et al., 2014). Such changes may underlie value modulation of early visual processing during VDAC (e.g., the aforementioned P1 effects), although in the current study we did not find evidence for such early value modulation. Increased decodability of reward-associated stimuli (e.g., Hickey and Peelen, 2015) has also been taken as supporting evidence (Failing and Theeuwes, 2018). Importantly, Hickey and Peelen (2015) were decoding object categories. In the current study, we focused on within-category stimulus identity and did not observe value modulation of identity decoding in either the reward learning or attention task. Moreover, in line with previous research on VDAC (e.g., Anderson et al., 2011; Hickey and Peelen, 2015), reward learning in the current study was conducted over a shorter timescale than perceptual learning studies, which typically require multi-day training (e.g., Schoups et al., 2001). It remains unclear whether visual cortical plasticity can occur under the same timescale as VDAC.

### Value signals outside the cortex

Alternatively to the visual cortical plasticity account, VDAC may be explained by plasticity in subcortical regions (Anderson, 2016; Hikosaka et al., 2018), to which MEG is relatively insensitive (Hillebrand and Barnes, 2002). One route may be a recently identified circuit involved in rapid orienting to reward-associated stimuli, which includes the tail of the caudate nucleus, the substantia nigra pars reticulata, and the superior colliculus (Griggs et al., 2018; Hikosaka et al., 2018; Yamamoto et al., 2012, 2013; Yasuda and Hikosaka, 2015). Learned value modulates neural activity in each of these regions, which can ultimately affect visual cortical processing via the thalamus (Hikosaka et al., 2018).

Value information can affect selective attention via multiple parallel pathways (Hikosaka et al., 2018). Models of cortico-basal ganglia interactions suggest that value may first be encoded in the basal ganglia, which gradually establishes stimulus-value associations in the cortex (Hélie et al., 2015; Houk and Wise, 1995). Consistent with this, value coding in LIP and extrastriate cortical neurons shifts earlier (e.g., earlier than 200ms) after days of training (Frankó et al., 2010; Peck et al., 2009). It is therefore possible that with longer training we would have observed earlier cortical value signals in the current study. Other candidate regions for reward-related plasticity include the amygdala and hippocampus, both of which show learning-driven value coding (Chau et al., 2015; Paton et al., 2006; Peck and Salzman, 2014; Wimmer et al., 2018).

### Conclusion

Overall, our results suggest that VDAC is underpinned by learned value signals which modulate spatial selection throughout posterior visual and parietal cortex. However, despite previous studies suggesting an important role for early cortical plasticity, we suggest that VDAC can occur in the absence of changes in early processing in cortex.

## Conflict of interest

The authors declare no competing financial interests.

## Acknowledgements

This research was funded by a Biotechnology and Biological Sciences Research Council (BB/M010732/1) and James S. McDonnell Foundation Scholar Award (220020405) to Mark G. Stokes, and by the NIHR Oxford Health Biomedical Research Centre. The Wellcome Centre for Integrative Neuroimaging is supported by core funding from the Wellcome Trust (203139/Z/16/Z), and The Netherlands Organisation for Scientific Research (NWO Veni grant 016.Veni.198.065 awarded to E.S.).

